# Synergy pattern of short cationic antimicrobial peptides against multidrug-resistant *Pseudomonas aeruginosa*

**DOI:** 10.1101/639286

**Authors:** Serge Ruden, Annika Rieder, Thomas Schwartz, Ralf Mikut, Kai Hilpert

## Abstract

With the rise of various multi-drug resistance pathogenic bacteria, worldwide health care is under pressure to respond. Conventional antibiotics are failing and the development of novel classes or alternative strategies is a major priority. Antimicrobial peptides (AMPs) can not only kill multi-drug resistant bacteria, but also can be used synergistically with conventional antibiotics. We selected 30 short AMPs from different origins and measured their synergy in combination with Polymyxin B, Piperacillin, Ceftazidime, Cefepime, Meropenem, Imipenem, Tetracycline, Erythromycin, Kanamycin, Tobramycin, Amikacin, Gentamycin, and Ciprofloxacin. In total 403 unique combinations were tested against a multi-drug resistant *Pseudomonas aeruginosa* isolate (PA910). As a measure of the synergistic effects, fractional inhibitory concentrations (FICs) were determined using microdilution assays with FICs ranges between 0.25 and 2. A high number of combinations between peptides and Polymyxin B, Erythromycin and Tetracycline were found to be synergistic. Novel variants of Indolicidin also showed a high frequency in synergist interaction.

## Introduction

Among the most serious problems health care is facing is the increased number of infections caused by antibiotic-resistant bacteria, which can no longer be treated with previously active antimicrobial agents. The World Health Organization (WHO) identified in 2013 the development of antibiotic resistance as one of the major global threats to human society and recommended intensive monitoring for the identification of critical hot spots, in order to reduce transmission. The global spread of antibiotic resistance is one of the most interesting examples of biological evolution. It is highly relevant to both human and animal health and welfare with a direct impact on society. The primary cause of this situation is the excessive use of antibiotics (Berendonk et al., 2015). Although the environmental bacteria are most probably the original source of many antibiotic resistance genes found in bacterial pathogens, the impact of nosocomial resistance and transmission has greatly increased over the last half century (Bush et al., 2011). The increased prevalence of antibiotic resistance in microbiota is due to four major reasons: i) horizontal transfer of antibiotic resistance genes, ii) the assortment of resistant bacteria due to selective pressures from antimicrobial usage, (iii) the bacterial ability for gene mutation and recombination (e.g. presence of mutator determinants) (Cantón and Morosini, 2011; Davies and Davies, 2010; Wright and Sutherland, 2007) and (iv) the dissemination of resistant organisms from human and animal commensals that have adapted to antibiotic treatment of the host. Importantly, we cannot exclude proliferation of antibiotic resistance due to the spread of resistant bacterial clones and their mobile genetic elements carrying antibiotic-resistance genes, this is due to spontaneous processes not necessarily linked with antibiotic resistance (Baquero et al., 2013; Kohanski et al., 2010; Seiler and Berendonk, 2012). Though the over-use of antibiotics may select for resistant populations, other biotic and abiotic factors including physicochemical conditions, pollution, induction of stress responses, bacterial adaptation and phenotypic heterogeneity among others, can enhance the effect of selective pressure. Evidence has shown that even in sub-inhibitory concentration antibiotics may still exert their impact on microbial community (Andersson and Hughes, 2014).

The review on antimicrobial resistance chaired by Jim O’Neill, initiated by the UK prime minister in 2014, published 2016 estimates that by 2050 more people (10 million) will die each year from infections than current number of people who die from cancer (https://amr-review.org/Publications.html). In order to maintain modern medical standards of care, it is a matter of urgency that novel antimicrobials are discovered and developed, particularly those with novel modes of action, which are less likely to suffer cross-resistance to existing drugs. The WHO published a priority list in 2017 (http://www.who.int/news-room/detail/27-02-2017-who-publishes-list-of-bacteria-for-which-new-antibiotics-are-urgently-needed) of bacteria that are especially problematic, in order to provide information and focus for drug development projects. In the highest category is carbapenem-resistant *Pseudomonas aeruginosa*.

*Pseudomonas aeruginosa* is a rod-shaped, Gram-negative bacterium, which is naturally found in soil and water and therefore well adapted to humid environments. It is a clinically important, opportunistic pathogen, which may cause pneumonia and bacteraemia in the elderly or immuno-compromised hosts. It is responsible for chronic, destructive lung disease in patients suffering from cystic fibrosis (Bhagirath et al., 2016). *P. aeruginosa* exhibits a higher intrinsic resistance to a number of antimicrobial agents compared to most other Gram-negative bacteria (Yoneda et al., 2005). Additionally, rapid development of resistance to previously effective antimicrobials, such as fluoroquinolones, sulfonamides or macrolides has been observed (Breidenstein et al., 2011; Ellington and Woodford, 2006; Oh et al., 2003; Sköld, 2000).

Unfortunately, there has been a significant reduction in the development of novel antimicrobial agents with many major pharmaceutical companies halting research in anti-infective agents. The fact there are very few new antimicrobial agents with a new mode of action increases the risk of a nightmare scenario where even “minor” infections could become serious health risks. As there is already only a limited number of anti-pseudomonal antibiotics and increasing prevalence of resistance, it is an important question as to whether potential new antibiotics with different modes of action also synergise with “old” antimicrobials, especially for multi-drug resistant (MDR) bacteria. Antimicrobial peptides (AMPs) are potential novel antimicrobial drugs with some much-desired features, for example low chance for development of resistance, fast acting, broad spectrum activity and active against MDR bacteria. So far only a few have been enrolled in clinical studies (Czaplewski et al., 2016; Greber and Dawgul, 2017). In this study we investigated whether synergy exists between designed short, cationic AMPs and conventional antibiotics in MDR *P. aeruginosa* or not. Such a synergy could be a starting point for further optimization to re-use already clinically valuable antibiotics.

## Material and Methods

### Peptides and antibiotics

All peptides used in this study were purchased from the Brain-Research Centre at the University of British Columbia. Peptides were characterized and purified via high-performance liquid chromatography (HPLC), mass was confirmed by matrix assisted laser desorption time of flight (MALDI-TOF) mass spectroscopy. Purity of all peptides was greater than 90%.

All antibiotics were purchased from VWR, except Polymyxin B, which was purchased from Sigma-Aldrich.

### Minimal inhibitory concentration (MIC)

We selected six different *P. aeruginosa* isolates that were described as multi-drug resistant and had been isolated from clinical or municipal waste water (Schwartz et al., 2006). The MIC was determined in a microdilution assay using Mueller-Hinton broth (MH) following a previous published protocol (Wiegand et al., 2008). Briefly, a twofold serial dilution of the antibiotics and peptides were prepared and added to a bacteria solution resulting in 2-5 10^5^ CFU/ml. The microtiter plates (polypropylene, Corning) were incubated for 18 h at 37 °C and MIC were taken visually.

### Fractional inhibitory concentration (FIC)

The checkerboard assay was used to determine the FICs, following the protocol described in Koneman’s Color Atlas and Textbook of Diagnostic Microbiology (Winn Jr et al., 2006). Briefly, combinations of peptides and antibiotics were prepared in 96 well plates (polypropylene, Corning) in a twofold dilution series. After the addition of a log-phase bacterial inoculum of 2-5 x 10^5^ CFU/ml, plates were incubated at 37 °C for 18 h. The FICs were determined by visual inspection. The effects of the combinations were determined using the FICs. The FIC was computed by adding two partial FIC values, FIC_A_ - the MIC of drug A, tested in combination with drug B divided by the MIC of drug A, tested alone and FIC_B_. - the MIC of drug B, tested in combination with drug A divided by the MIC of drug B, tested alone (FIC=FIC_A_+FIC_B_= (MIC_AB_/MIC_A_) + (MIC_BA_/MIC_B_), where MIC_A_ and MIC_B_ are the MICs of drugs A and B alone, respectively, and MIC_AB_ and MIC_BA_ are the MIC concentrations of the drugs in combination). Here we use the European Committee for Antimicrobial Susceptibility Testing (EUCAST) definition, which is very similar to the definition provided by Odds 2003, except Odds define additive effects when FIC values are between 0.5 and 1 (European Committee for Antimicrobial Susceptibility Testing (EUCAST) of the European Society of Clinical Microbiology and Infectious Dieases (ESCMID), 2000; Odds, 2003). The combination of peptide (drug A) and antibiotics (drug B) was defined as synergistic if the FIC was ≤0.5, indifferent if the FIC was >0.5 but ≤4.0 and antagonistic if the FIC was >4.

## Results

First, we have determined the MIC values against a range of different antibiotics for all six strains of *P. aeruginosa*, that were described as multi-drug resistant and had been isolated from clinical or municipal waste water, in comparison with a sensitive wild type strain (PAO1), see Table 1 (Schwartz et al., 2006). This confirmed that these strains are multi-drug resistant.

**Table 1.** MIC determination measured at least in triplicates (n=3) against wild type P. aeruginosa and MDR isolates (values in µM)

| Antibiotic/Strain | PA01-wt | PA 910 | PA 915 | PA 919 | PA 923 | PA 927 | PA253 |
| --- | --- | --- | --- | --- | --- | --- | --- |
| Polymyxin B | 0.125 | 0.125 | 0.25 | 0.125 | 0.25 | 0.25 | 0.125 |
| Ciprofloxacin | 0.25 | 64 | 32 | 64 | 64 | 16 | 64 |
| Tobramycin | 0.25-0.5 | 32 | 64 | 64 | 32 | 32 | 64 |
| Gentamycin | 1 | 128 | 128-256 | 128 | 128 | 128 | >256 |
| Amikacin | 1 | 2-4 | 4 | 32 | 4 | 4 | 8-16 |
| Imipenem | 2 | > 128 | >128 | 16 | > 128 | >128 | > 128 |
| Meropenem | 2-4 | >128 | >128 | >128 | >128 | >128 | >128 |
| Piperacillin | 4 | 64-128 | >256 | 64 | 128 | 256 | >256 |
| Ceftazidime | 4 | 128 | 4 | 8 | 128 | >128 | > 128 |
| Tetracyclin | 8 | 64 | 16 | 32 | >512 | 16 | 32 |
| Cefepime | 8 | 64 | 32 | 16 | 64 | 128 | >128 |
| Kanamycin | 64 | 128-256 | 256-512 | 128 | 256 | 128 | >512 |
| Erythromycin | 128 | 128 | 64 | 128 | 256 | 16 | 64 |

A set of short peptides between 9 and 13 amino acids in length were selected from already published and from ongoing unpublished work (Cherkasov et al., 2009; Hilpert et al., 2005, 2006; Mikut et al., 2016), LL37 was used as a comparison to long and helical peptides. The MIC values of the peptides against the wild type *P. aeruginosa* strain as well as three selected MDR variants of PA910, PA919 and PA253 were determined (Table 2). The majority of MIC values were equal or similar (plus/minus factor of 2) between the wild type and the MDR strains, with the highest change observed for Indolicidin with a 8-fold decrease in activity. Since the wt PA01 shows in some cases a higher sensitivity towards the antimicrobial peptides we tested with three peptides if this relates to the MDR phenotype or the adaption to a specific environment. We tested 25 clinical isolates with full sensitivity and 25 clinical isolates with various resistance and MDR pattern (data not shown). Nearly identical MIC variations were determined in both groups, demonstrating that the observed sensitivity of the wt strain is not related to the missing expression/carriage of resistant genes.

**Table 2.**
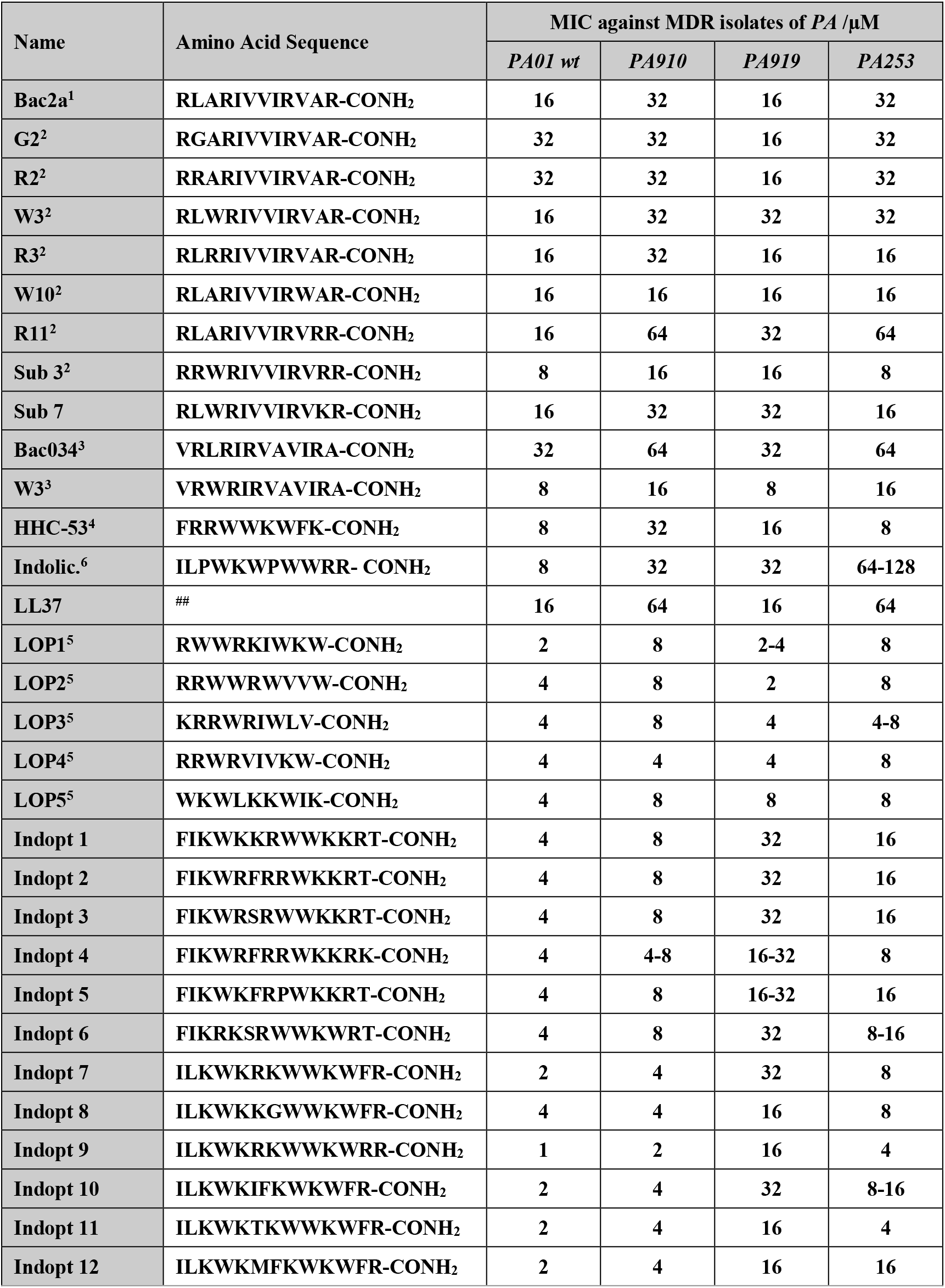
MIC values in µM of short AMPs (9-13mer) measured in triplicates (n=3) against several isolates of MDR P. aeruginosa. MIC values of all Indopt variants were measured in 1/8 MHb, whilst all other were measured in full MHb.. A 37mer human AMP was included as compassion (LL37). ## The sequence of the peptide LL-37: LLGDFFRKSKEKIGKEFKRIVQRIKDFLRNLVPRTES-COOH. ^1^ (Wu and Hancock, 1999), ^2^ (Hilpert et al., 2005), ^3^ (Hilpert et al., 2006), ^4^ (Cherkasov et al., 2009), ^5^ (Mikut et al., 2016), ^6^ (Selsted et al., 1992)

Strain PA 910, which is only sensitive to Polymyxin B and Amikacin was selected to perform a synergy study combining 31 antimicrobial peptides with 12 antibiotics and 1 lipopeptide (Polymyxin B) resulting in 403 unique combinations.

The synergistic effects of combination of short antimicrobial peptides and conventional antibiotics resulted in a complex pattern. There were peptides that show no synergistic effect, whilst others showed a variety, similar to the tested antibiotics, see Figure 1A and B.

**Figure 1:**
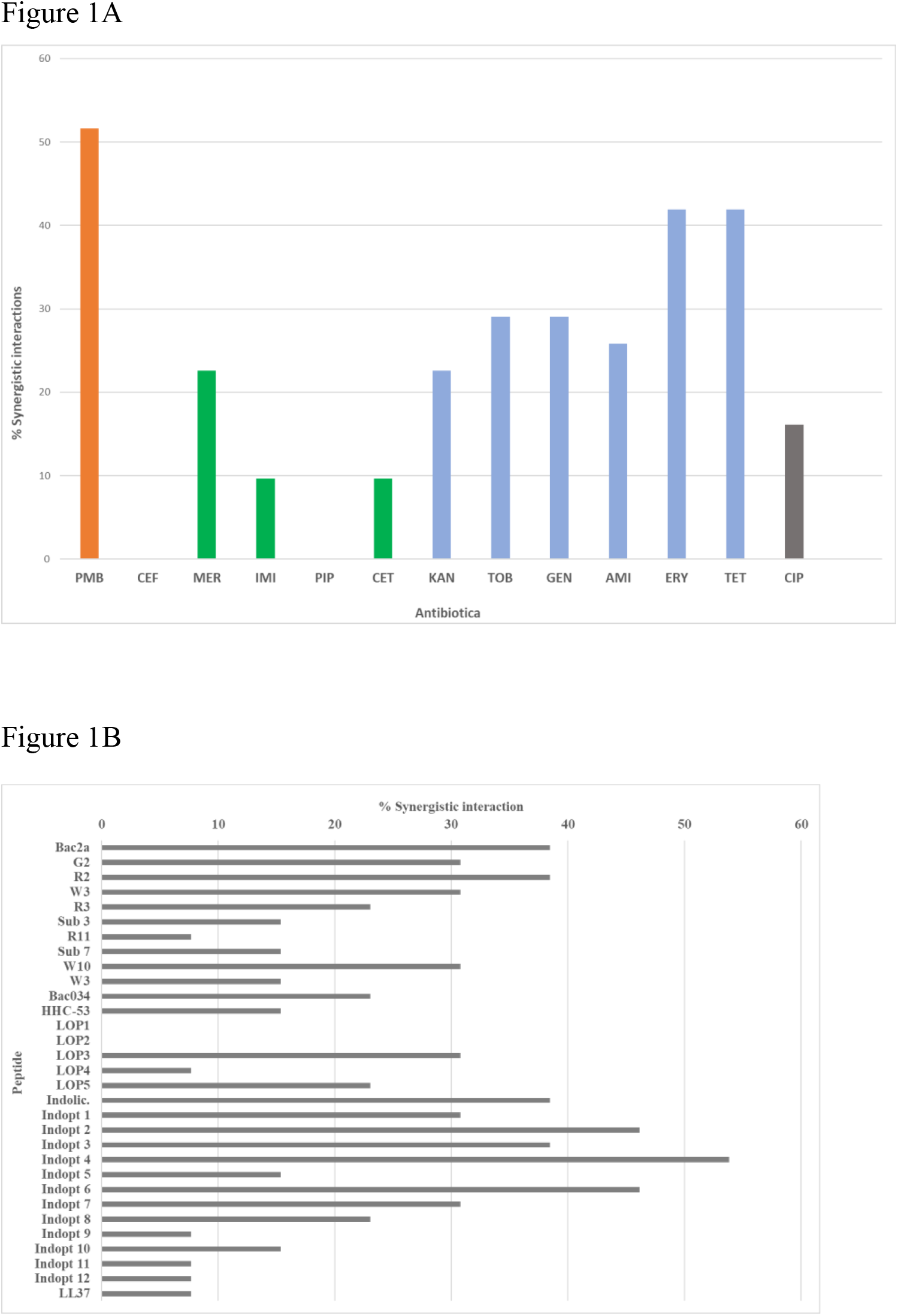
The maximal number of possible interactions was set to 100% and the percent synergistic interaction was calculated for A) conventional antibiotics and B) short antimicrobial peptides. Colour codes and abbreviations see Table 3

Selected peptide-antibiotic combinations were tested three times to confirm first findings, see Table 4.

**Table 3:**
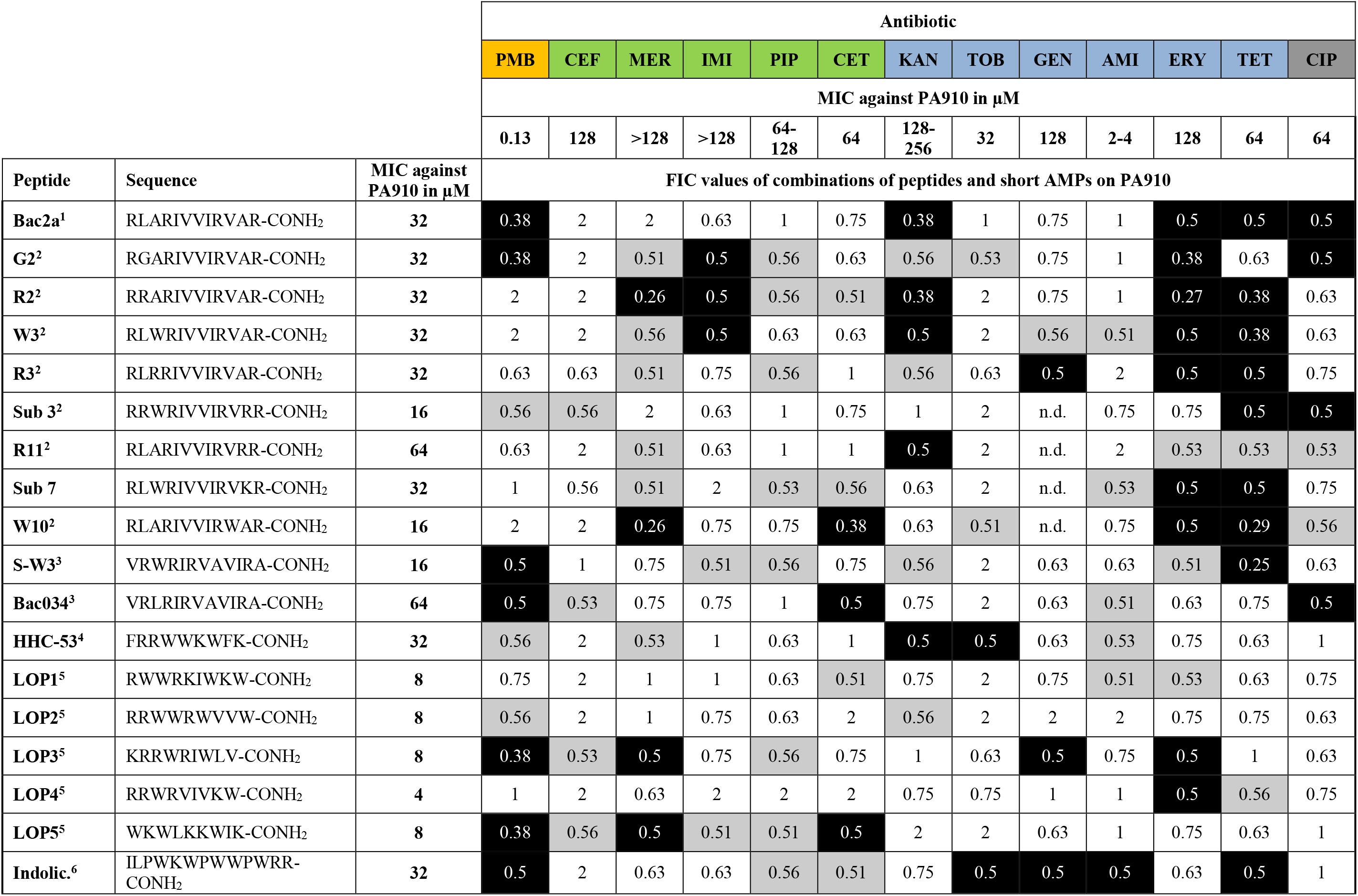
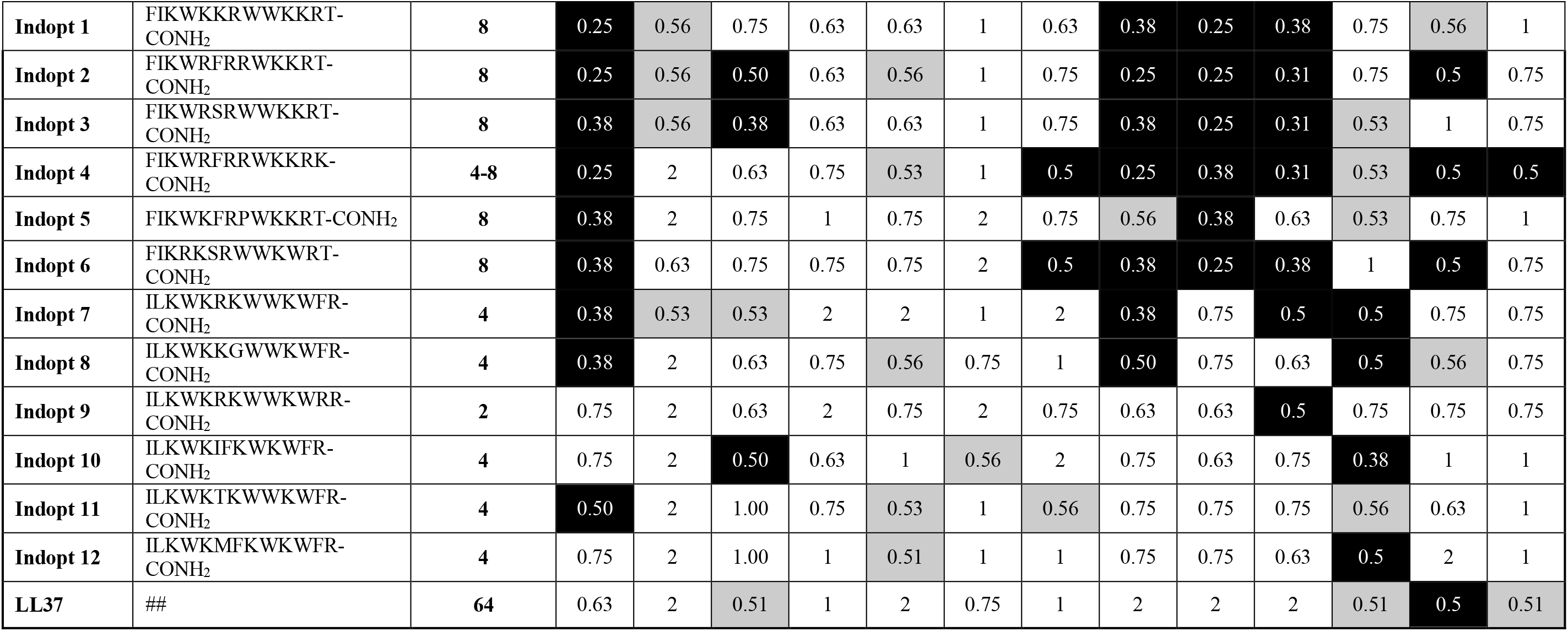
Fractional inhibitory concentrations (FICs) of short AMPs, 9-13mers, and different antibiotic were determined once (n=1) against an MDR isolate of P. aeruginosa (PA 910). All FICs were determined once using MHb, for the Indopt versions, 1/8 MHb was used. FIC values equal or smaller than 0.5 are considered as synergistically interaction and highlighted in white colour on black background. FIC values equal or smaller than 0.6 (a 20% error margin) are considered as potentially synergistic and were labelled in grey. PMB was used for the lipopeptide Polymyxin B, which is acting on cell wall and membrane, colour code in orange. Piperacillin (PIP) is of the ureidopenicillin class, Ceftazidime (CET) third-generation cephalosporin, Cefepime (CEF) fourth-generation cephalosporin, Meropenem (MER) and Imipenem (IMI) are part of the carbapenem subgroup. They all act on the cell wall, especially cell wall synthesis, colour coded in green. Tetracycline (TET), Erythromycin (ERY), Kanamycin (KAN), Tobramycin (TOB), Amikacin (AMI) and Gentamycin (GEN) are all acting at different sites of the bacterial ribosomes, colour coded in light blue. Ciprofloxacin (CIP) is a gyrase inhibitor abs acts on the DNA replication system, colour coded in grey. ## The sequence of the peptide LL-37: LLGDFFRKSKEKIGKEFKRIVQRIKDFLRNLVPRTES-COOH, n.d. stands for not determined in this test.

**Table 4:** Mean values of at least three measurements (n=3) and standard deviation, values in brackets, of fractional inhibitory concentrations (FICs) of selected short AMPs, 9-13mers, and different antibiotic were determined against an MDR isolate of P. aeruginosa (PA 910). FICs were determined in MHb, for the Indopt versions, 1/8 MHb was used. FIC values equal or smaller than 0.5 are considered as synergistically interaction and highlighted in white colour on black background. FIC values equal to or smaller than 0.6 (a 20% error margin) are considered as potentially synergistic and were labelled in grey. Colour codes and abbreviations see Table 3. All peptides that showed FIC values equal or smaller than 0.5 in the first experiment (see Table 3) that were not included in this confirmation study are labelled with n.d..

| Peptide | Antibiotic |  |  |  |  |  |  |  |  |  |  |  |  |
| --- | --- | --- | --- | --- | --- | --- | --- | --- | --- | --- | --- | --- | --- |
|  | PMB | CEF | MER | IMI | PIP | CET | KAN | TOB | GEN | AMI | ERY | TET | CIP |
| Bac2a<sup>1</sup> | 0.47<br>(0.09) |  |  |  |  |  | 0.63<br>(0.17) |  |  |  | 0.69<br>(0.21) | 0.44<br>(0.08) | 1.08<br>(0.61) |
| G2<sup>2</sup> | 0.45<br>(0.16) |  |  | 1.00<br>(0.70) |  |  |  |  |  |  | 0.63<br>(0.25) | 0.53<br>(0.01) | 0.59<br>(0.11) |
| R2<sup>2</sup> | 1.09<br>(0.79) |  |  | 1.00<br>(0.70) |  |  | 0.63<br>(0.17) |  |  |  | 0.42<br>(0.10) | 0.38<br>(0.10) |  |
| W3<sup>2</sup> | 1.17<br>(0.72) |  |  | n.d. |  |  | 0.67<br>(0.11) |  |  |  | 0.67<br>(0.11) | 0.38<br>(0.25) |  |
| R3<sup>2</sup> | 0.46<br>(0.14) |  |  |  |  |  |  | n.d. | 0.67<br>(0.11) |  | 1.04<br>(0.64) | 0.46<br>(0.05) |  |
| Sub 3<sup>2</sup> |  |  |  |  |  |  |  |  |  |  |  | 0.50<br>(0.00) | n.d. |
| R11<sup>2</sup> | 0.42<br>(0.14) |  |  |  |  |  | 0.75<br>(0.17) |  |  |  |  | 0.43<br>(0.12) |  |
| Sub 7 | 0.67<br>(0.31) |  |  |  |  |  |  |  |  |  | n.d. | 0.46<br>(0.06) |  |
| W10<sup>2</sup> |  |  | n.d. |  |  | n.d. |  |  |  |  | n.d. | 0.60<br>(0.27) |  |
| S-W3<sup>3</sup> | 0.81<br>(0.59) |  |  |  |  |  |  |  |  |  | 0.46<br>(0.11) | 0.34<br>(0.06) |  |
| Bac034<sup>3</sup> | 0.43<br>(0.12) | 0.59<br>(0.11) | 0.58<br>(0.11) |  |  | 0.50<br>(0.00) |  |  |  |  |  | 0.50<br>(0.17) | 0.75<br>(0.17) |
| HHC-53<sup>4</sup> | 0.46<br>(0.10) |  |  |  |  |  | 0.60<br>(0.10) |  | 0.54<br>(0.06) |  |  | 0.52<br>(0.10) |  |
| LOP1<sup>5</sup> | 0.55<br>(0.18) |  |  |  |  |  |  |  |  |  |  | 0.55<br>(0.19) |  |
| LOP2<sup>5</sup> | 0.60<br>(0.13) |  |  |  |  |  |  |  |  |  |  |  |  |
| LOP3<sup>5</sup> | 0.40<br>(0.04) |  | 0.75<br>(0.17) |  |  |  |  |  | 0.50<br>(0.00) |  | 0.44<br>(0.08) | 0.71<br>(0.19) |  |
| LOP4<sup>5</sup> |  |  |  |  |  |  |  |  |  |  | n.d. |  |  |
| LOP5<sup>5</sup> | 0.38<br>(0.00) |  | 0.54<br>(0.06) |  |  | n.d. |  |  |  |  | 0.50<br>(0.17) | 0.50<br>(0.10) |  |
| Indolic.<sup>6</sup> | n.d. |  |  |  |  | 0.71<br>(0.21) |  | n.d. |  | n.d. |  | n.d. |  |
| Indopt 1 | 0.35<br>(0.10) |  | 0.59<br>(0.14) |  |  | 0.59<br>(0.29) |  | 0.38<br>(0.00) | 0.21<br>(0.03) | 0.38<br>(0.00) | 0.48<br>(0.18) |  |  |
| Indopt 2 | 0.38<br>(0.08) |  | 0.54<br>(0.06) |  |  |  |  | 0.29<br>(0.03) | 0.19<br>(0.04) | 0.32<br>(0.04) |  | 0.75<br>(0.17) |  |
| Indopt 3 | 0.33<br>(0.06) |  | 0.38<br>(0.00) |  |  |  |  | 0.33<br>(0.06) | 0.23<br>(0.10) | 0.37<br>(0.08) | 0.47<br>(0.06) |  |  |
| Indopt 4 | 0.29<br>(0.06) |  | 0.50<br>(0.09) |  |  |  | 0.60<br>(0.10) | 0.27<br>(0.07) | 0.23<br>(0.03) | 0.31<br>(0.04) |  | 0.59<br>(0.10) | 0.63<br>(0.25) |
| Indopt 5 | 0.33<br>(0.06) |  | 0.47<br>(0.19) |  |  |  |  | 0.52<br>(0.03) | 0.35<br>(0.03) | 0.55<br>(0.05) |  |  |  |
| Indopt 6 | 0.38<br>(0.00) |  | 0.52<br>(0.15) |  |  |  | 0.55<br>(0.06) | 0.38<br>(0.08) | 0.27<br>(0.07) | 0.38<br>(0.00) |  | 0.50<br>(0.00) |  |
| Indopt 7 | 0.38<br>(0.00) |  |  |  |  |  |  | 0.67<br>(0.22) |  | 0.55<br>(0.05) | 0.50<br>(0.00) |  |  |
| Indopt 8 | 0.29<br>(0.06) |  | 0.54<br>(0.06) |  |  |  |  | 0.58<br>(0.11) |  |  | 0.45<br>(0.07) |  |  |
| Indopt 9 | 0.54<br>(0.14) |  |  |  |  |  |  | 0.46<br>(0.14) |  | 0.51<br>(0.01) |  |  |  |
| Indopt 10 | 0.46<br>(0.19) |  | 0.52<br>(0.03) |  |  |  |  | 0.63<br>(0.17) |  |  | 0.38<br>(0.00) |  |  |
| Indopt 11 | 0.46<br>(0.06) |  |  |  |  |  |  | 0.52<br>(0.15) |  |  |  |  |  |
| Indopt 12 | 0.48<br>(0.18) |  |  |  |  |  |  |  |  |  | 0.59<br>(0.06) |  |  |
| LL37 |  |  |  |  |  |  |  |  |  |  |  | n.d. |  |

To test whether the observed synergy was dependent on the selected strain or not, two additional MDR PA isolates (PA253 and PA919) were used to determine the FICs of the selected peptides used with Polymyxin B, see Table 5.

**Table 5.**
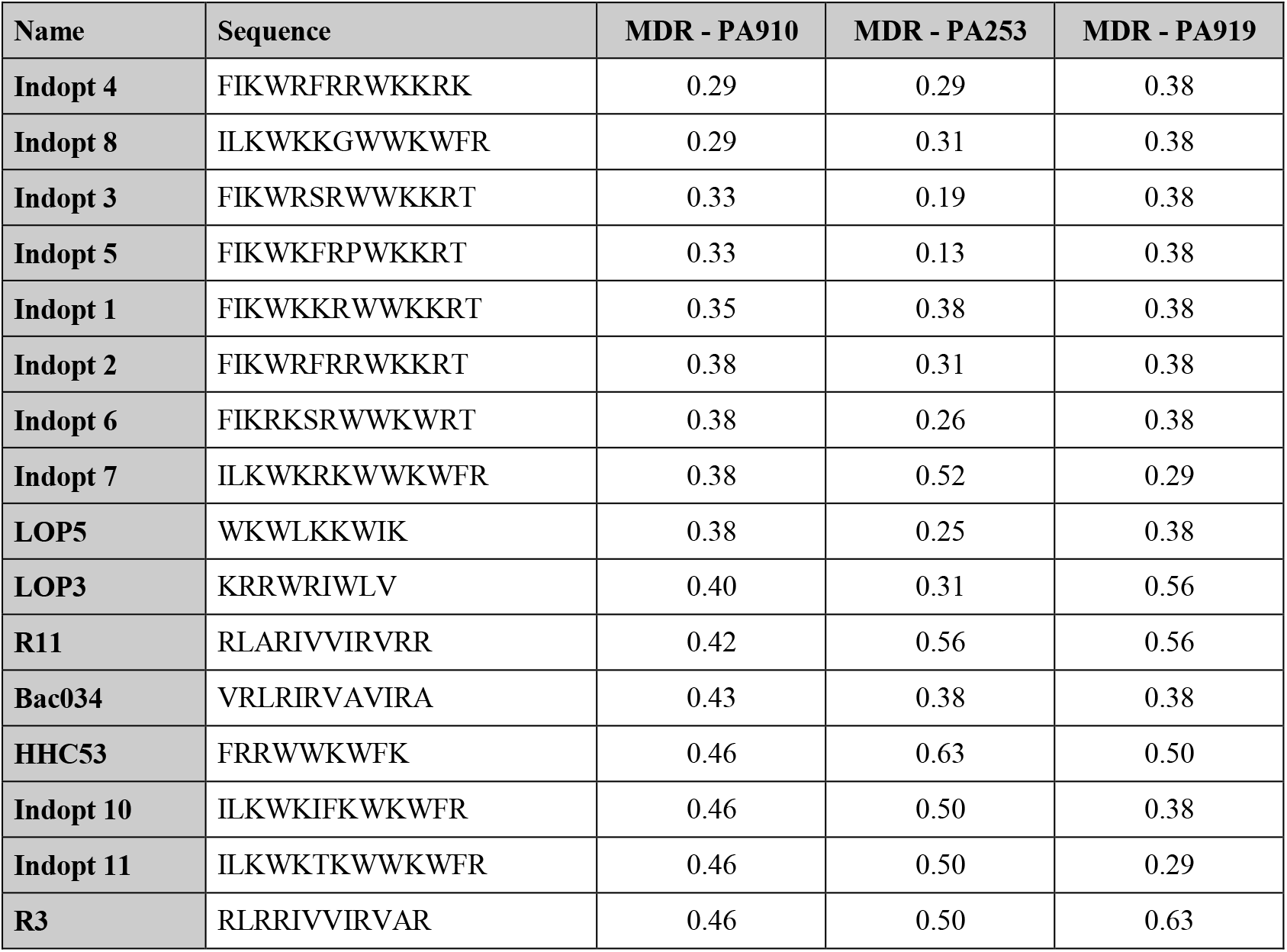
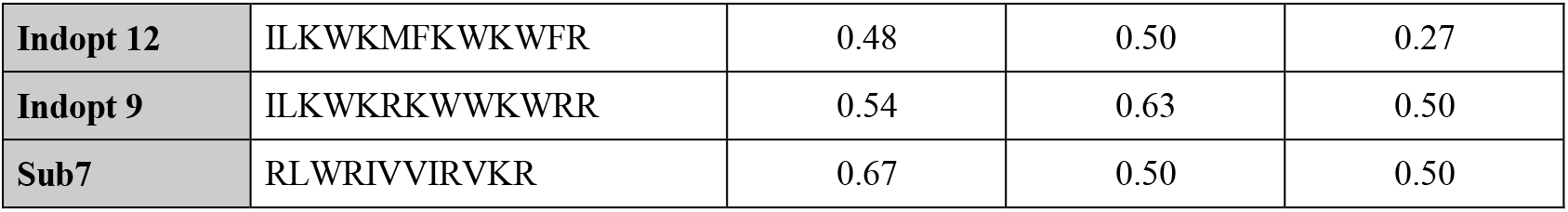
FIC values of selected peptides for three different MDR P. aeruginosa strains against Polymyxin B. Values from MDR-PA910 are mean values from at least three different measurements (n=3), values from MDR-PA253 and MDR-PA919 are measured once (n=1). Table is sorted according the values from MDR-PA910, from low to high.

## Discussion

*P. aeruginosa* is ranked amongst the top five organisms causing bloodstream, urinary tract, pulmonary, surgical site and soft tissue infections in patient in intensive care units (Veesenmeyer et al., 2009). The bacterium is widely distributed in the environment, as it can utilize a wide range of materials for its nutrients, whilst only requiring a limited amount of nutrients to survive (Abdelraouf et al., 2011). The current treatment regimen for multi-drug resistant cases being limited to the last resort antibiotic colistin (Hachem et al., 2007; Sabuda et al., 2008). However, alarmingly there have been reports of *P. aeruginosa* resistance to colistin (Goli et al., 2016). The situation is getting very concerning, in fact the World Health Organization has declared it to be a “critical priority pathogen,” on which research and development of novel antibiotics should be focused (Tacconelli et al., 2018).

We confirmed the described multi-drug resistance of *P. aeruginosa* strains that were isolated in clinical and municipal waste water, see Table 1 (Schwartz et al., 2006). It has been reported that antimicrobial peptides (AMPs) show synergy with conventional antibiotics both in planktonic and biofilm growth (Giacometti et al., 2000; Jorge et al., 2017). Here we studied if a set of short antimicrobial peptides that we have developed in previous projects can synergize with antibiotics in order to revive them. We also included Polymyxin B, since treatment failures with monotherapy of Polymyxins are reportedly increasing making it an urgent candidate for use with synergistic agents. There are currently clinical trials investigating Colistin alone versus Colistin in combination with Meropenem (ClinicalTrials.gov IDs NCT01732250 and NCT01597973) (Lenhard et al., 2016).

The FICs between combinations of antibiotics and peptides show several enhanced synergistic effects by using antibiotics in tandem with short AMPs. In general, the beta-lactams and beta-lactam like antibiotics show the lowest amount of synergy, along with Ciprofloxacin, a gyrase inhibitor, which also showed a rather low amount of synergy. Cefepime and Piperacillin showed no detectable synergy at all. Whereas, antibiotics acting on the ribosome show a higher amount of synergy, with the highest proportion of synergy observed with Polymyxin B, which acts on the cell wall and cell membrane. The majority of the results were confirmed by three independent measurements of the selected combinations. Some combinations however showed a larger FIC values as determined in the first screen, this proving the importance of verification of FIC data (Hsieh et al., 1993).

Polymyxin B binds to the lipid A portion of the lipopolysaccharides (LPS), to replace cationic ions like Ca^2+^ and Mg^2+^ from the LPS layer (Morrison and Jacobs, 1976). This process destabilizes the LPS layer, leading to permeability changes and consequently to an “self-promoted uptake” (Hancock, 1997; Hermsen et al., 2003). This process destabilises the membrane and allows molecules to pass through the membrane in both directions. It has been shown that Polymyxin B synergises with antibiotics (Abdul Rahim et al., 2015; Elemam et al., 2010; Zusman et al., 2013) as well with antimicrobial peptides (Draper et al., 2013; Giacometti et al., 2000; van der Linden et al., 2009). The majority of short cationic peptides used in these studies also showed synergistic interaction with Polymyxin B (Table 4). This synergy was verified for two additional strains (Table 5). Small changes in the sequence of Bac2A leads to a loss in synergy, this is especially pronounced by the introduction of a tryptophan residue, for example W3, Sub3, W10 and S-W3. The data suggests that an additional tryptophan residue might anchor the peptide more in the membrane and consequently does not support synergy with Polymyxin B.

Cefepime is a fourth-generation cephalosporin antibiotic that has an extended spectrum of activity against Gram-positive and Gram-negative bacteria and is more stable as compared to third-generation agents (Wynd and Paladino, 1996). Cefepime has good activity against multi-drug resistant *Streptococcus pneumoniae*. Cephalosporins are bactericidal that disrupts the synthesis of the peptidoglycan layer of the bacterial cell walls by blocking transpeptidases known as penicillin binding proteins (PBPs) (Klein and Cunha, 1995). There was no synergistic effect observed, indicating that the peptides did not improved access to target sites or that they interfered with the lactamases. Ceftazidime is a third-generation cephalosporin antibiotic targeting PBPs. Similar to Cefepime there were no synergistic combinations determined with the exception of Bac034, W10 and LOP5. Meropenem, a carbapenem type beta-lactam antibiotic active against Gram-positive and Gram-negative bacteria, blocks PBPs and shows a bactericidal activity (Blumer, 1997). In contrast to Cefepime, a range of short AMPs showed synergistic interaction. Imipenem on the other hand, another carbapenem type beta-lactam with the same mode of action, did show three synergistic interactions; however, in the confirmation experiment (Table 4) these were not verified (Park and Parker, 1986). We conclude that synergy is caused by a Meropenem-specific feature, for example increasing the uptake rate for this molecule. Piperacillin is a penicillin beta-lactam antibiotic used mainly for gram-positive organisms. Piperacillin demonstrates bactericidal activity as a result of the inhibition of cell wall synthesis by binding to PBPs. Piperacillin is stable against hydrolysis by a variety of beta-lactamases, including penicillinases, cephalosporinases and extended spectrum beta-lactamases (Eliopoulos and Moellering, 1982). No synergistic combinations were observed for Piperacillin with the short AMPs tested.

Kanamycin (Kanamycin A) belongs to the aminoglycoside class and is a natural compound found in *Streptomyces kanamyceticus* (de Lima Procópio et al., 2012). Aminoglycosides bind to the 30S subunit of the ribosome: most binding occurs on the 16 srRNA of the bacteria leading to a bactericidal action (Walter et al., 1999). It shows broad-spectrum activity against Gram-negative bacteria and some activity against Gram-positive. From all the aminoglycosides, Kanamycin shows the least activity and the least confirmed synergy. Tobramycin, is produced in *Streptomyces tenebrarius* and belongs to the class of aminoglycoside antibiotics. It binds irreversible to the 30S ribosomal subunit and shows a broad-spectrum activity, especially effective against *P. aeruginosa* (https://www.drugbank.ca/drugs/DB00684). Gentamicin is an aminoglycoside antibiotic that is produced by *Micromonospora purpurea* and acts on the 30S subunit. It is highly active against Gram-negative bacteria and shows activity against some Gram-positive bacteria (Daniels et al., 1975). Amikacin is a semi-synthetic antibiotic based on Kanamycin A, both members of the class aminoglycosides, which binds to the 30S subunit and 16 srRNA. Amikacin demonstrates a broad-spectrum activity towards Gram-negative bacteria, including pseudomonades and has some effects on Gram-positive bacteria, including *Staphylococcus aureus* (Ristuccia and Cunha, 1985). The synergy patterns of Tobramycin, Gentamycin and Amikacin are very similar to the synergy pattern of Indolicidin, especially to the variants of Indopt 1-8. This is in accordance to research published by Boehr at al., showing that Indolicidin and analogues (differ from the ones in this study) are able to inhibit aminoglycoside phosphotransferase and aminoglycosides acetyltransferase. Through this inhibition the most effective resistance mechanism can be weakened (Boehr et al., 2003). Based on these findings a further optimization of Indopt peptides could lead to the formation of more potent inhibitors to overcome aminoglycosides resistance.

Erythromycin is a macrolide antibiotic produced by *Saccharopolyspora erythraea* and reversibly binds to the 50S subunit of the bacterial ribosome (https://www.drugbank.ca/drugs/DB00199). It is active against Gram-negative and Gram-positive bacteria. Tetracycline is a natural produced antibiotic by *Streptomyces aureofaciens* and binds reversibly to the 30S subunit as well as to some extent also the 50S subunit, with potential influence over the bacterial membrane (https://www.drugbank.ca/drugs/DB00759). Tetracycline belongs to the class of tetracyclines and is a broad-spectrum antibiotic with activity against Gram-positive and Gram-negative bacteria. The synergy patterns for these two classes look different from the aminoglycosides. There is a broader range of peptides that can synergise with these antibiotics for example Bac2A variants, 9mer variants and the indolicidin variants.

Ciprofloxacin is a synthetic antibiotic belonging to the fluoroquinolones and is an inhibitor of the bacterial topoisomerase II (DNA gyrase) and topoisomerase IV. Ciprofloxacin is a broad-spectrum antibiotic with a wide range of Gram-positive and Gram-negative bacteria (https://www.drugbank.ca/drugs/DB00199). There are only few peptides showing synergy, all at 0.5 and confirmation studies showed even higher values for the selected combinations.

In summary, our data shows that small antimicrobial peptides that can kill multi-drug resistant bacteria, in this case MDR *Pseudomonas aeruginosa*, are able to synergise with conventional antibiotics despite the fact that they are not effective anymore. We believe that this shows the potential to develop these molecules not only as mono-therapeutic agents, but also as part of a combination therapy with the conventional antibiotics to re-use antibiotics by this synergistic approach with AMPs. This can be an alternative avenue for dealing with the resistance crisis.

